# Biofabrication of Small Vascular Graft with Acellular Human Amniotic Membrane: A Proof-of-Concept Study in Pig

**DOI:** 10.1101/2024.09.11.612466

**Authors:** O Aung, Peter J Rossi, Mitchell R Dyer, Austin Stellpflug, Yingnan Zhai, Allen Kenneth, Xiaolong Wang, Jackie Chang, Yiliang Chen, Brandon Tefft, Rongxue Wu, Lingxia Gu, Bo Wang

## Abstract

Synthetic vascular grafts, such as expanded polytetrafluoroethylene (ePTFE), are commonly used for large vessel surgeries [internal diameter (ID) ≥ 10 mm] but present significant challenges in medium to small vessels (ID < 10 mm) due to increased risks of thrombosis, stenosis, and infection. In this study, we developed a small-diameter vascular graft using decellularized human amniotic membrane (DAM graft) (ID = 6 mm) and transplanted it into porcine carotid arteries, comparing it with ePTFE grafts to assess inflammation, biocompatibility, patency, and overall function. One-week post-implantation, ultrasound imaging confirmed blood patency in both graft types. However, after one-month, gross examination revealed pronounced neointimal hyperplasia in ePTFE grafts, while DAM grafts maintained open lumens without signs of stenosis or thrombosis. Histological analysis showed extensive fibrous tissue formation in ePTFE grafts, resulting in luminal narrowing, whereas DAM grafts displayed sustained lumen patency and vascular integration. Immunofluorescence confirmed reduced inflammation and improved tissue organization in DAM grafts, characterized by lower macrophage infiltration and better cellular architecture. These findings suggest that DAM grafts offer superior biocompatibility and significantly lower risks of neointimal hyperplasia, making them a promising alternative for small-diameter vascular surgeries compared to ePTFE grafts.

## 1. Introduction

Vascular diseases, including cerebrovascular disease, coronary heart disease, aneurysms, deep vein thrombosis, and peripheral arterial disease, pose serious health risks due to their high morbidity and mortality rates and require a multifaceted management approach [1, 2]. This includes lifestyle modifications, pharmacotherapy, endovascular procedures, and surgical interventions [2-4].

While lifestyle changes are crucial, they are often insufficient in advanced cases. Pharmacological treatment encompasses the use of many medications such as antiplatelet agents (e.g., aspirin), statins (to mitigate cholesterol levels), antihypertensive drugs, and anticoagulants (e.g., warfarin or direct oral anticoagulants) to manage symptoms and mitigate underlying risk factors associated with vascular pathologies [6, 7]. With the development of medical devices and surgical techniques, intravascular stents have been developed, offering minimally invasive endovascular procedures aimed at alleviating narrowed or obstructed blood vessels, thereby enhancing blood circulation [8-10]. However, stents can still face issues like acute thrombosis and restenosis, which may affect long-term effectiveness [12-16]. In more severe cases, surgical options such as bypass grafting or vascular replacement are necessary to restore blood flow by using functional blood vessel substitutes [11].

Autologous blood vessels, such as internal mammary artery and radial artery harvested from patients themselves, are always regarded as the gold standard vascular substitute for vascular surgeries. The major advantages of autologous blood vessels include their inherent biocompatibility, low risk of thrombosis, and favorable long-term durability [17-20]. However, the availability of suitable autologous vessels is limited in some patients with pre-existing chronic diseases or previous vessel harvest, and the associated morbidities of vessel harvesting, and subsequent surgeries further restrict their therapeutic applicability [21, 22].

Synthetic grafts, including polytetrafluoroethylene (ePTFE), polyethylene terephthalate (PET), and polyurethane (PU), have received approval from the U.S. Food and Drug Administration (FDA) for clinical application [23-28]. These materials exhibit good mechanical and biological compatibility with native blood vessels and have shown positive results in large vessel surgeries [internal diameter (ID) > 6mm]. However, clinical trials targeting medium (ID: 6-10 mm) to small (ID < 6mm) vessels, particularly in coronary arteries, infrainguinal arteries (below the inguinal ligament), and infrageniculate arteries (below the knee), often result in unsatisfactory outcomes. This is largely due to factors such as decreased blood flow velocity and increased resistance in smaller vessels, which lead to higher rates of thrombosis, intimal hyperplasia, stenosis, occlusion, and infection [29, 30].

In our previous study, we developed a small-diameter vascular graft using decellularized human amniotic membrane (DAM) with an ID = 1.3 mm [31]. The DAM graft was created by rolling DAM around a catheter, followed by lyophilization and crosslinking with 1% genipin. This produced a multilayered structure with mechanical properties similar to native arteries. *In vitro* studies showed good cellular attachment, migration, and compatibility with human endothelial cells. *In vivo* testing, where a portion of the rat abdominal aorta was replaced without anticoagulants, demonstrated long-term patency for over 16 months. The graft supported endothelial and smooth muscle regeneration and, critically, showed no signs of neointimal hyperplasia. These results indicate that the DAM graft holds strong potential for future small-diameter vascular reconstruction surgeries.

Although the findings from our previous study are promising, the DAM grafts used in rat surgeries are much smaller than those required for human vascular procedures. Clinically, small-diameter vascular grafts are needed for surgeries involving peripheral arteries (ID ≈ 5 mm) [32], coronary arteries (ID = 3-5 mm) [33], and for hemodialysis access in end-stage renal disease patients (ID ≈ 6 mm) [34]. The aim of this study is to develop a DAM graft that meets the structural, mechanical, and biological requirements of human vascular reconstruction. To assess its clinical potential, we will implant the DAM graft into the porcine carotid artery and compare its performance with that of ePTFE grafts, focusing on evaluating the inflammatory response, biocompatibility, patency, and overall functionality of the DAM graft *in vivo*.

## 2. Materials and Methods

### 2.1. Fabrication of DAM graft

To match the size of the porcine carotid artery, DAM vascular grafts with an inner diameter (ID) of 6 mm were fabricated. Fresh human amniotic membranes were sourced from the tissue bank at the Medical College of Wisconsin (MCW) and decellularized using a solution of 1% Triton X-100 and 0.1% sodium dodecyl sulfate over approximately five days. The decellularization process was followed by thorough water washing to prepare the DAM [35-37].

To ensure the DAM grafts had suitable mechanical properties for carotid artery replacement, we prepared three variations with different numbers of layers in the DAM wall. Rectangular DAM pieces measuring 20 cm × 8 cm, 25 cm × 8 cm, and 30 cm × 8 cm were rolled around a 6 mm diameter catheter. During this process, a 1% elastin solution was applied to enhance adhesion between the layers. Following rolling, the DAM grafts underwent complete lyophilization using a Freeze-Dryer System (Cole-Parmer, Vernon Hills, IL) at -80°C. Subsequently, they were crosslinked using a 0.5% w/v genipin solution for 24 hours at room temperature, followed by thorough rinsing with water. The crosslinked DAM grafts were then stored at 4°C until further experimentation [31].

### 2.2. Histology and SEM Characterizations

For histology staining, DAM grafts were fixed in 10% formalin, dehydrated through a series of ethanol concentrations, and cleared with xylene. The specimens were embedded in paraffin and sectioned into 6 µm thick slices for hematoxylin and eosin (H&E) staining. The cross-sectional areas of the grafts were then examined using light microscopy.

For scanning electron microscopy (SEM) analysis, the grafts were fixed with 2.5% glutaraldehyde for 24 hours, dehydrated using a graded ethanol series, and subjected to critical point drying. Following this, the specimens were sputter-coated with gold-palladium and the cross-sectional area of the grafts was observed using SEM (LEO Gemini 1525).

### 2.3. Mechanical testing

The uniaxial tensile properties of ePTFE and DAM grafts with varying layers (n=4 per group) were tested using a TA rheology machine (TA Instruments, New Castle, DE). Each sample was stretched at a rate of 0.1 mm/s until failure. Stress was calculated by normalizing the applied force to the initial cross-sectional area, while strain was determined by dividing the displacement by the initial grip-to-grip distance (gauge length at 1 g preload).

### 2.4. Transplantation into porcine carotid artery

Based on histological and mechanical evaluations, we chose the DAM graft with a 15-layer wall structure for in vivo testing. This graft was implanted into both carotid arteries of two pigs to evaluate blood compatibility and graft patency. As controls, we implanted ePTFE grafts with an ID = 6 mm (Becton, Dickinson & Company) into the carotid arteries of two additional pigs and used healthy porcine carotid arteries as a positive control in two more pigs.

Pigs weighing between 18-23 kg, both males and females, were procured from Wilson’s Prairie View Farm, Inc. (Burlington, WI). Prior to surgery, the pigs underwent a 24-hour fasting period, followed by post-operative fluid feeding for one day and subsequent normal feeding for one month. Anesthesia was induced with an intramuscular injection of atropine, tiletamine/zolazepam, and xylazine, and maintained using 1-5% isoflurane with 100% O2 inhalation through an anesthetic machine.

The ventral neck region was shaved using clippers, and the surgical site was prepared by disinfecting with betadine followed by alcohol wipes repeated thrice. Each pig underwent bilateral grafting, with identical vascular grafts implanted into both the left and right carotid arteries. This standardized approach facilitated comparative analysis between the two graft sites and minimized variability in the experimental setup. Pigs received local anesthesia by intramuscular injection of Telazol (25 mg/1 kg body weight) and Xylazine (20 mg/1 kg body weight). Pigs also inhaled general anesthesia by isoflurane (1-5%). Heparin (300 mg/1 kg of body weight) was administered intravenously prior to graft transplantation to prevent clot formation. The carotid arteries were cross-clamped, and Papaverine was applied as needed for vasodilation. Protamine sulfate (1.3 mg/1 kg of body weight) was administered intravenously after graft implantation to prevent heparin overdose. A segment of the carotid artery (approximately 2-3 cm long) was excised, and either the DAM graft or ePTFE graft (5-6 cm in length) was anastomosed to the remaining carotid artery using an end-to-end technique with 8-0 nylon sutures. The cross-clamps were then removed to restore blood flow. The incisions were closed in three layers using absorbable sutures (3-0 vicryl) to close the strap muscles, subcutaneous tissue, and skin, with Dermabond used to protect the incision site.

### 2.5. Ultrasound imaging

After 1 week of transplantation surgery, the patency of the implanted grafts was evaluated by the ultrasound imaging (Echocardiography Core at MCW) utilizing a GE Vivid IQ ultrasound system equipped with an L8-18i transducer (GE Healthcare, WI, USA). All the pigs for ultrasound imaging test underwent anesthesia using the method described in section 2.4. Standard 2-D imaging with color Doppler was employed to visualize the carotid artery and the implanted vascular graft. Pulsed Wave Doppler analysis was utilized to measure the velocity of blood flow within the implanted graft. Additionally, ultrasound imaging was performed on pigs that did not undergo vascular surgery to serve as a comparison for native artery characteristics.

### Histological and immunofluorescent staining of grafts

One month after the surgery, all the pigs were anesthetized, and the implanted grafts in DAM and ePTFE groups or the carotid arteries in the control group were harvested and fixed in 4% formaldehyde. For histology staining, all the segments were processed using the protocol outlined in Section 2.2, and stained with H&E and sirius red/fast green kits (Sigma-Aldrich, St. Louis, MO) following with the manufacturer protocols. For immunofluorescence staining, slides were stained with alpha-smooth muscle actin (α-SMA), CD31, and CD68 primary antibodies at 4°C overnight. The secondary antibodies goat anti-rabbit IgG H&L (Alexa Fluor® 488) and donkey anti-rat IgG H&L (Alexa Fluor™ Plus 594) (abcam, Cambridge, MA) were applied at room temperature for 1 hour. Negative controls were prepared using the same protocol without the primary antibody.

### 2.7. Statistical Analysis

All data presented were expressed as the mean ± standard deviation. Comparisons were determined by Analysis of variance (ANOVA SAS 9.0). The Holm–Sidak Test was used for post-hoc comparisons. Data correlations were determined from mean values using a Pearson-product moment correlation (parametric) or Spearman Rank Order Correlation (nonparametric). Results were considered significantly different at *p* < 0.05.

## 3. Results

### 3.1. Morphological characterization of DAM grafts

The DAM vascular grafts, fabricated from different-sized DAM sheets, consistently maintained tubular structures with varying wall thicknesses and layering. Specifically, the grafts made from 20 cm × 8 cm DAM sheets contained approximately 10 layers in the vascular wall (Fig. 1A). Those created from 25 cm × 8 cm DAM sheets featured about 15 layers (Fig. 1B), while the grafts made from 30 cm × 8 cm DAM sheets had approximately 20 layers (Fig. 1C). These results, confirmed by H&E staining and SEM imaging, also showed that all DAM grafts had a tightly adhered, multi-layered wall structure.

**Figure 1.**
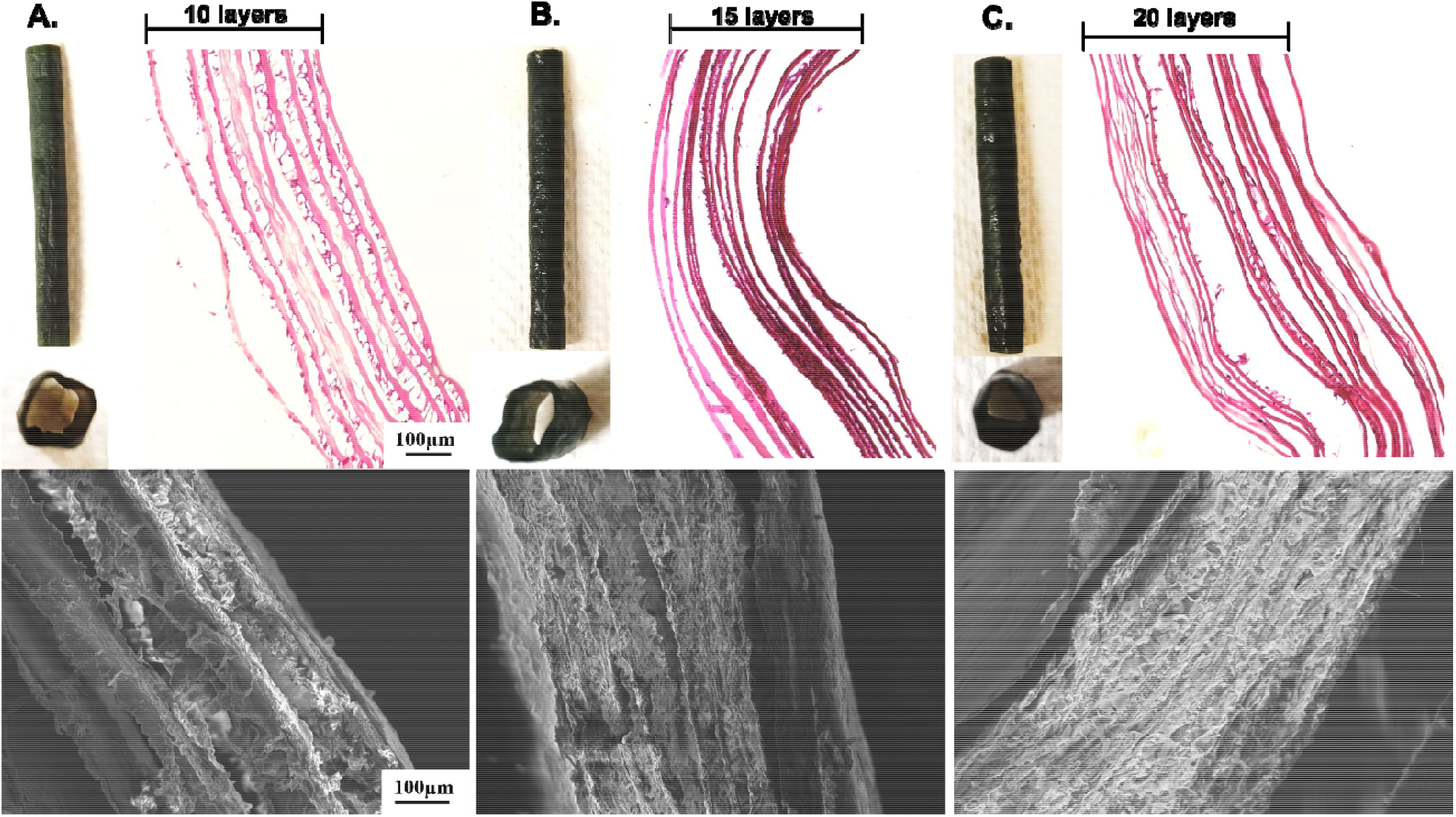
Longitudinal view (top left), H&E staining (top right), and SEM imaging (bottom) of (A) 10-layer DAM graft; (B) 15-layer DAM graft; and (C) 20-layer DAM graft.

### 3.2. Mechanical characterization of DAM vascular grafts

The uniaxial tensile testing and the tensile modulus comparison revealed that DAM grafts at 10, 15 and 20 layers were statistically greater than ePTFE graft (DAM graft 10 layers: 5652.3 +-748.2 kPA; DAM graft 15 layers: 4845.6 +-550 kPa; DAM graft 20 layers: 4664.8 +-559.6 kPa; ePTFE graft: 3225 +-272.3 kPA; (DAM 10 vs ePTFE: (p=0.006); DAM 15 vs ePTFE: (p= 0.01); DAM 20 vs ePTFE: (p = 0.01) via ANOVA) (Fig. 2 A, B). The tensile modulus of DAM grafts at 10 ,15 and 20 layers showed no statical significance when compared to each other (DAM 10 vs DAM 15: (p=0.20); DAM 10 vs DAM 20: (p= 0.14); DAM 15 vs DAM 20: (p = 0.71) via ANOVA) (Fig. 2 A-B).

**Figure 2.**
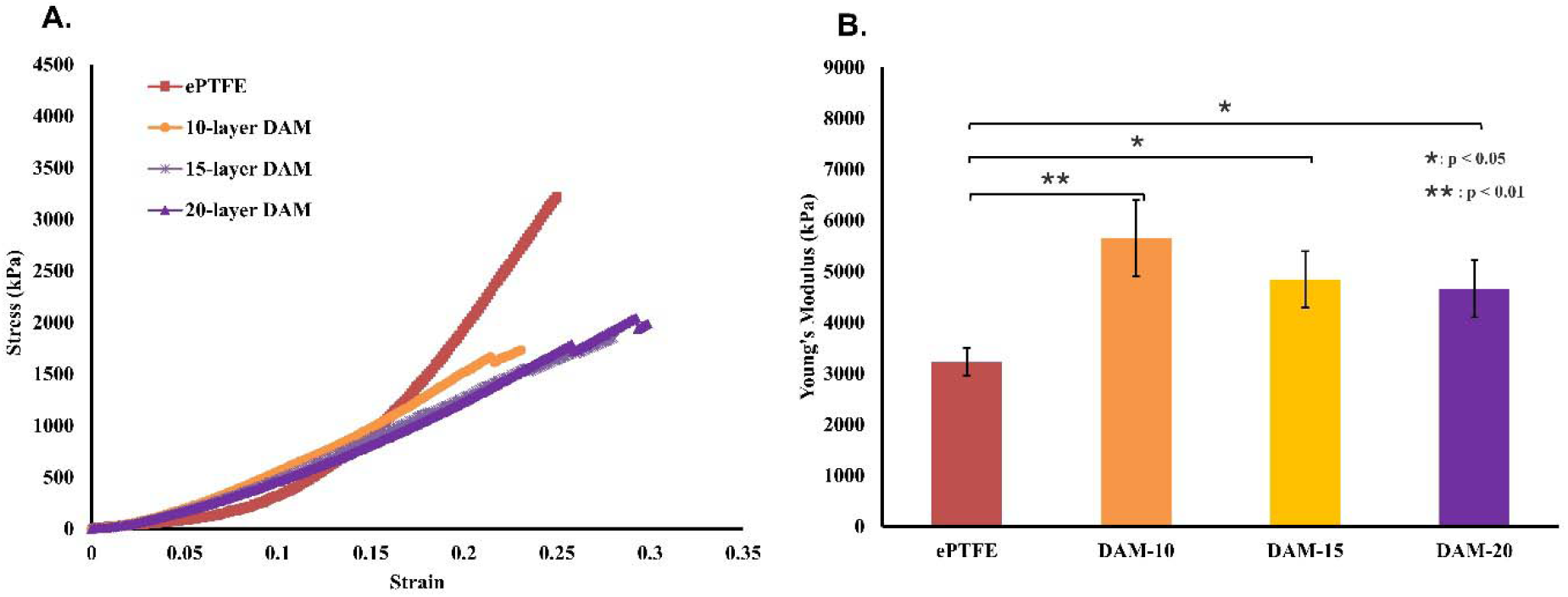
(A) Uniaxial tensile property and (B) Young’s Modulus of ePTFE, and varying layers of DAM grafts. ^*^ indicates p<0.05, ^**^ indicates p<0.01.

### 3.3. Graft patency in porcine carotid artery

During transplantation surgery, both the DAM (Fig. 3B) and ePTFE (Fig. 3C) grafts were successfully sutured without issues such as tearing, bleeding, or oozing. Postoperatively, all animals showed normal behavior and activity with no signs of infection or complications. 1-week post-transplantation, ultrasound imaging demonstrated patent grafts with blood flow in both the DAM (Fig. 3E) and ePTFE (Fig. 3F) groups, confirming the sustained functionality of the implanted grafts.

**Figure 3.**
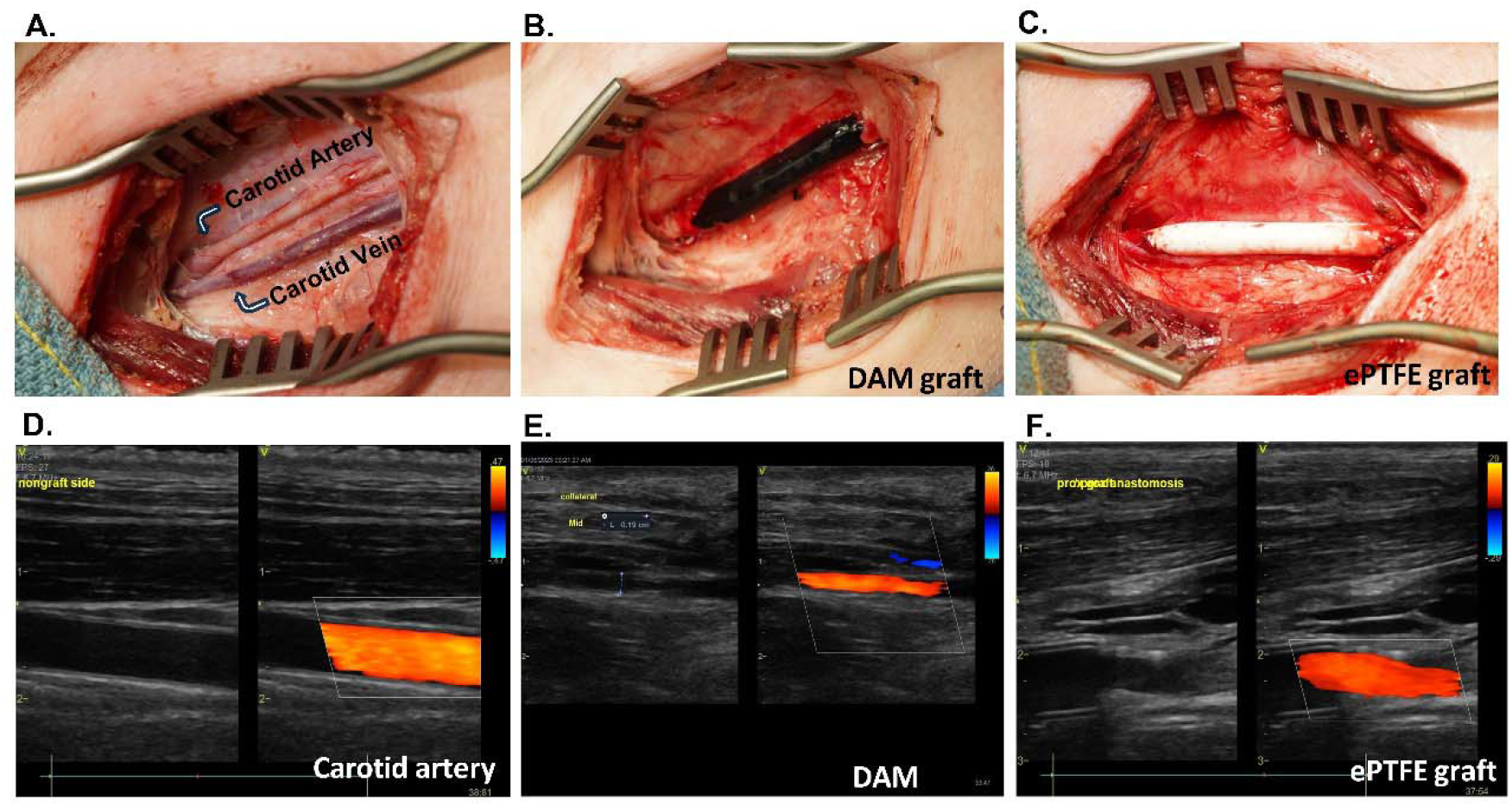
(A) Native carotid artery of pig; (B) DAM graft surgically implanted in a pig carotid artery (C) ePTFE graft surgically implanted in a pig carotid artery; (D) Ultrasound image of native carotid artery; (E) Ultrasound image of DAM graft 1 week after vascular transplantation; (F) Ultrasound image of ePTFE graft 1 week after vascular transplantation.

### 3.4. Gross view of the implanted grafts

After 1-month of transplantation surgery, ePTFE and DAM grafts, along with normal carotid arteries from the control group, were harvested for further evaluation. Gross examination revealed significant and uniform neointimal hyperplasia within the lumen of the ePTFE grafts (Fig. 4A, C), originating from the anastomosis site compared to the original ePTFE graft (Fig. 4B). In contrast, DAM grafts (Fig. 4A, E) exhibited minimal neointimal hyperplasia compared to the original DAM graft (Fig. 4D), demonstrating superior performance in this regard.

**Figure 4.**
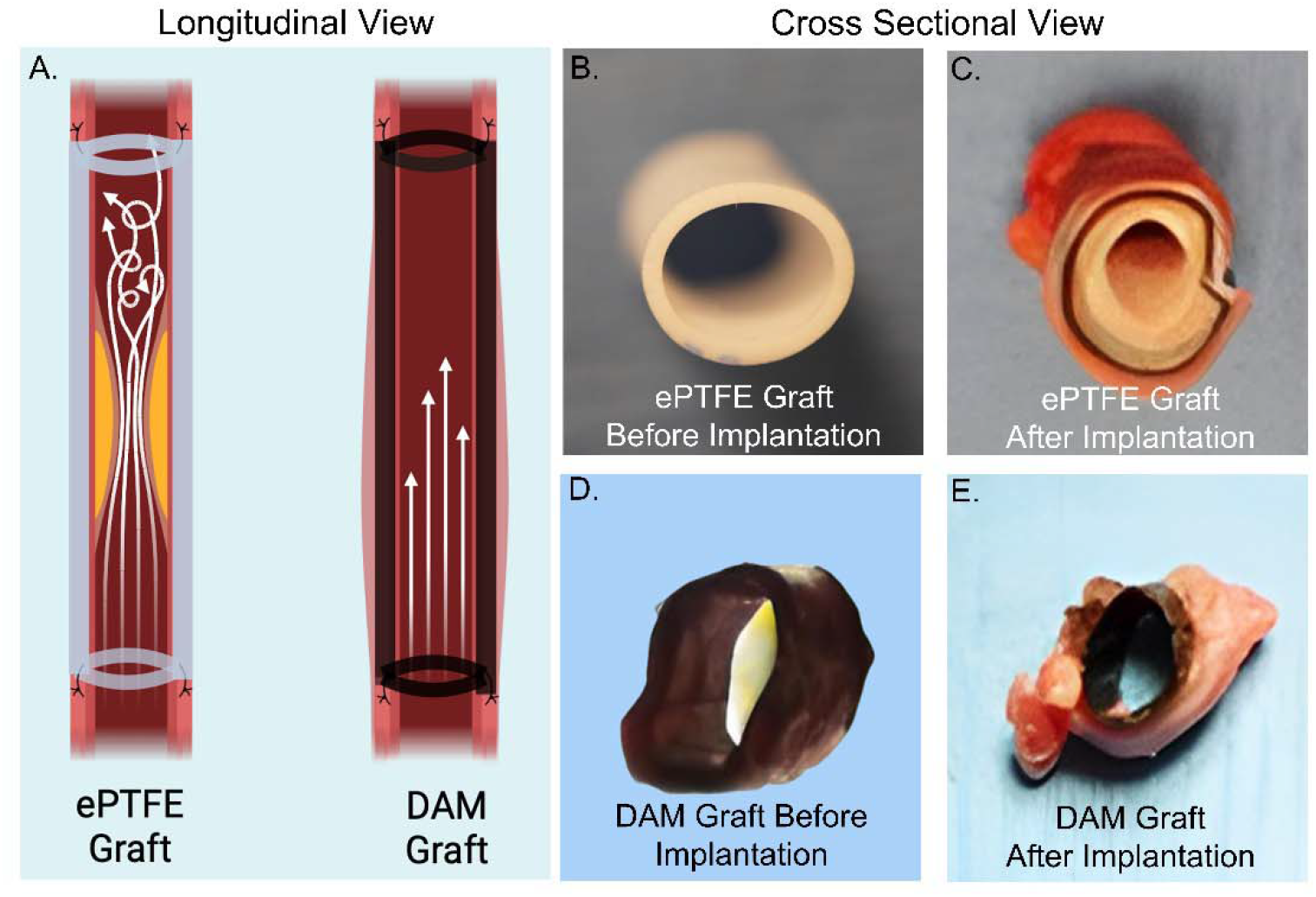
(A) Longitudinal model of ePTFE and DAM grafts 1 month after surgical transplantation; (B) Cross sectional view of ePTFE graft before surgical transplantation; (C) Cross sectional view of ePTFE graft 1 month after surgical transplantation; (D) Cross sectional view of DAM graft before surgical transplantation; (E) Cross sectional view of ePTFE graft 1 month after surgical transplantation.

### 3.5. Histology and immunofluorescence staining

Hematoxylin and eosin (H&E) staining revealed distinct differences in tissue response between the two graft types. Compared to the native pig artery (Fig. 5A), The ePTFE grafts demonstrated a thick layer of fibrous tissue overlaying the lumen, predominantly composed of stellate cells surrounded by connective tissue (Fig. 5 I). This fibrous layer contributed to the luminal narrowing observed in the ePTFE grafts. Conversely, the DAM grafts exhibited cellular invasion primarily from the outer layer of the graft, with the lumen surface remaining clear and devoid of stenosis, thrombosis, or deformation (Fig. 5E) compared to native pig artery (Fig. 5A). This indicates a more favorable integration and healing response with the DAM grafts compared to the ePTFE grafts.

**Figure 5.**
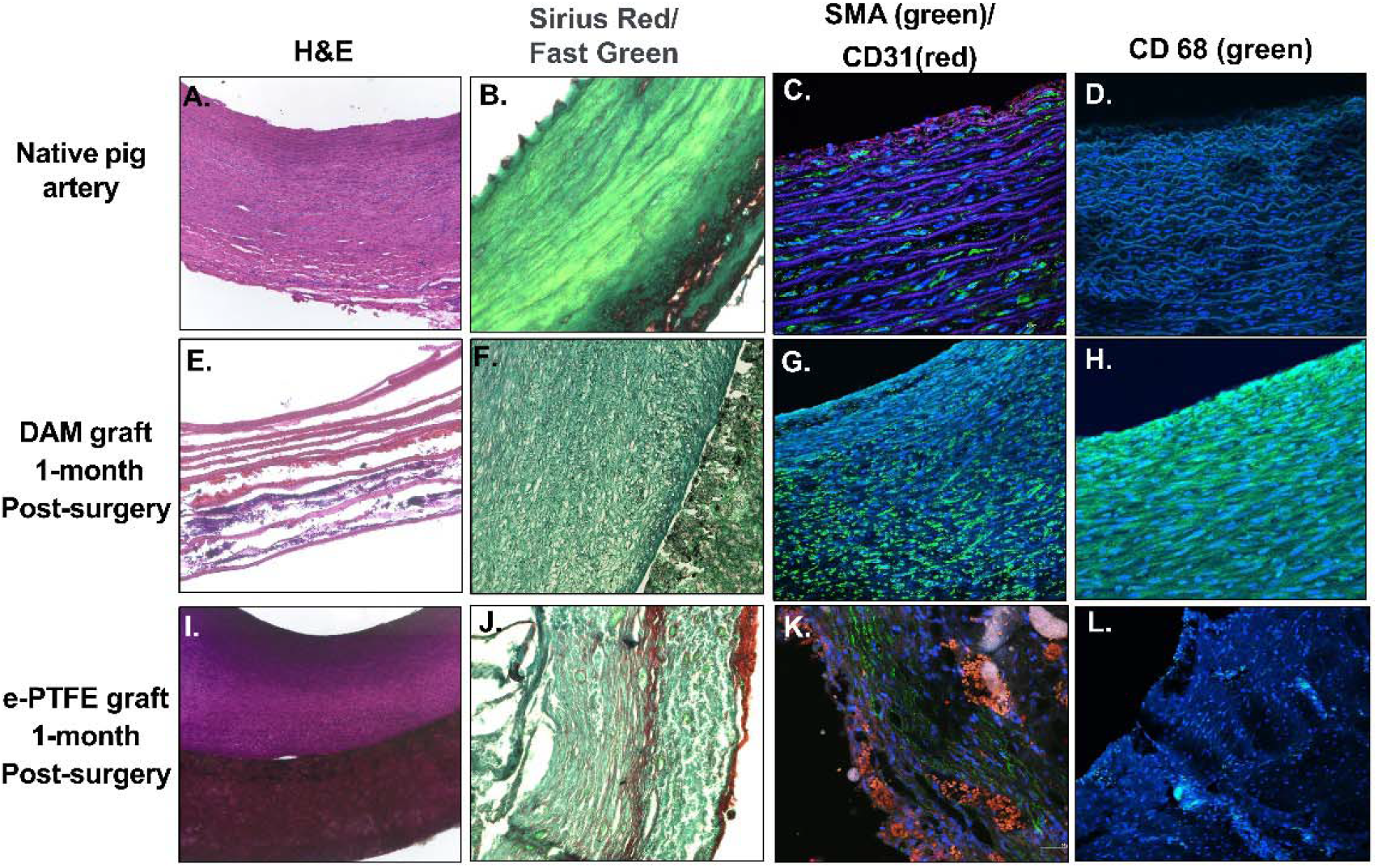
(A) H&E staining of native pig artery; (B) Immunofluorescence staining of native pig artery; (C) VE-cadherin a-SMA and CD31 staining of native pig artery; (D) CD 68 staining of native pig artery; (E) H&E staining of DAM graft 1 month after surgical implantation; (F) Immunofluorescence staining of DAM graft 1 month after surgical implantation; (G) VE-cadherin a-SMA and CD31 staining of DAM graft 1 month after surgical implantation; (H) CD 68 staining of DAM graft 1 month after surgical implantation; (I) H&E staining of ePTFE graft 1 month after surgical implantation; (J) Immunofluorescence staining of ePTFE graft 1 month after surgical implantation; (K) VE-cadherin a-SMA and CD31 staining of ePTFE graft 1 month after surgical implantation; (L) CD 68 staining of ePTFE graft 1 month after surgical implantation

Immunofluorescence staining was employed to characterize the cellular composition and organization within the ePTFE and DAM grafts one month post-implantation.

In the ePTFE grafts, immunofluorescent staining for smooth muscle actin (SMA) revealed a substantial presence of smooth muscle cells (SMCs) within the neointimal hyperplasia (Fig. 5J) compared to native pig artery (Fig. 5B). These SMCs were randomly organized, indicating disorganized cellular proliferation. Staining for CD31, an endothelial cell marker, showed an absence of endothelial cells in the neointimal layer, suggesting a lack of endothelialization in the ePTFE grafts (Fig. 5K) compared to native pig artery (Fig. 5C). Additionally, a significant number of CD68-positive macrophages were detected throughout the neointimal hyperplasia (Fig. 5L) compared to native pig artery (Fig. 5D). The abundance of macrophages indicates an ongoing inflammatory response, highlighting the immune-mediated remodeling processes in the ePTFE grafts.

In contrast, SMA staining identified a certain number of smooth muscle cells, which displayed a clear circumferential alignment, resembling the organized structure of the middle layer of a native vascular graft (Fig. 5F) compared to native pig artery (Fig. 5B). Furthermore, CD31 staining identified cells positive for CD31 around the luminal surface of the DAM grafts (Fig. 5G) compared to native pig artery (Fig. 5C). These cells, however, lacked nuclei, indicating they were red blood cells rather than endothelial cells. This suggests that while there was some interaction with blood components, endothelialization of the luminal surface had not occurred in the DAM grafts. The DAM grafts exhibited a markedly different cellular response. CD68 staining revealed only a limited presence of macrophages, suggesting a reduced inflammatory response compared to the ePTFE grafts (Fig. 5H) compared to native pig artery (Fig. 5D).

## 4. Discussion

Our earlier investigations have demonstrated the successful application of DAM vascular grafts in rat models, highlighting their potential as promising alternatives for vascular surgeries [31]. However, the small size of the DAM grafts used in these rat surgeries (ID = 1.5 mm) limits their applicability for human vascular procedures, which typically require grafts with an ID of 4-6 mm.

To address this limitation, the present study aims to develop DAM vascular prostheses with larger ID suitable for small vascular surgeries in humans. We evaluated the mechanical properties in relation to the layered structure of the DAM grafts and *in vivo* vascular patency and function of the DAM grafts in large animal models. This research seeks to bridge the gap between the promising outcomes observed in rat studies and the practical requirements of human vascular procedures.

After decellularization, the DAM scaffold is devoid of cellular components while preserving the ECM structure and mechanical stiffness, as demonstrated in our previous studies [35, 37, 38]. One of the primary advantages of the DAM scaffold is its adaptability. Its thin, fragile, and flexible nature allows fabrication into grafts of various dimensions and mechanical properties, tailored to meet the specific requirements of different clinical scenarios. This adaptability is crucial for addressing the diverse needs of human patients, providing a versatile solution for a range of vascular applications [31].

In our study, we prepared DAM vascular grafts based on the dimensions of the porcine carotid artery, maintaining a consistent ID of 5 mm. We varied the number of DAM layers in the wall structure, creating grafts with 10, 15, and 20 layers. This variation allowed us to investigate the correlation between the layered structure and mechanical properties of DAM vascular grafts. Mechanical testing revealed a significant increase in stress for DAM grafts at 10, 15, and 20 layers compared to the ePTFE grafts, suggesting that the DAM graft.

The ability to customize the layered structure and mechanical properties of DAM vascular grafts offers significant advantages for tailoring grafts to individual patient requirements. For example, in high-pressure arterial environments where greater mechanical strength is necessary, grafts with more layers can be used. Conversely, in situations where flexibility and compliance are more critical, fewer layers can be employed. This customization potential enhances the clinical applicability of DAM scaffolds, making them a promising option for a wide range of vascular repair and reconstruction procedures. Moreover, the successful fabrication of DAM vascular grafts with varying mechanical properties highlights their potential in personalized medicine. By adjusting the layer composition, we can create grafts that match the specific biomechanical environment of the patient’s vasculature. This tailored approach improves the likelihood of successful integration and function, thereby increasing the overall effectiveness of the treatment.

To further evaluate the blood compatibility, graft patency, and vessel function of DAM vascular grafts, we selected a 15-layered DAM graft for in vivo transplantation into the carotid artery of a porcine model. This choice was based on its vascular wall thickness and mechanical properties, which closely match those of the native porcine carotid artery, making it an ideal candidate for comparison with the widely used synthetic ePTFE graft in clinic.

Both DAM and ePTFE grafts were successfully implanted during surgery, showing no signs of tearing, bleeding, or oozing. Postoperative observations revealed that all pigs exhibited normal behavior and activity, with no signs of infection or other complications. Ultrasound imaging at 1-week and 1-month post-transplantation confirmed the patency of both graft types, demonstrating adequate blood flow. This initial success in maintaining functional vascular conduits is promising for the potential clinical viability of DAM grafts.

One month after transplantation, the implanted grafts were retrieved and subjected to gross examination, revealing significant differences between the DAM and ePTFE grafts. The ePTFE grafts exhibited notable neointimal hyperplasia originating from the anastomosis sites, resulting in significant luminal narrowing. In contrast, the DAM grafts showed a clear and open lumen without any neointimal hyperplasia.

Histological staining including H&E and Sirius Red/Fast Green staining further emphasized these distinctions. The ePTFE grafts displayed a thick fibrous tissue layer primarily composed of stellate cells and connective tissue, contributing to luminal stenosis. Conversely, the DAM grafts exhibited cellular invasion predominantly from the outer layers, maintaining a lumen free from stenosis, thrombosis, or deformation. These findings as shown in Figure 4, suggests a more favorable healing response with the DAM grafts, facilitating better integration and preserving luminal patency compared to ePTFE grafts.

To gain a comprehensive understanding of the cellular composition and structure within the neointimal hyperplasia observed in the ePTFE grafts, we conducted immunofluorescence staining. Our findings revealed significant proliferation of smooth muscle cells (SMCs) within the neointimal layer, characterized by a disorganized cellular arrangement. This SMC proliferation is a typical response in synthetic vascular grafts and can lead to luminal narrowing and eventual graft failure. Importantly, we observed an absence of CD31-positive endothelial cells within the neointimal layer of the ePTFE grafts. Endothelial cells play a critical role in maintaining vascular health by preventing thrombosis and promoting vascular function. The lack of endothelialization on the graft surface suggests that it may not support adequate attachment and growth of endothelial cells, posing a risk for thrombotic complications and compromising long-term graft performance. Furthermore, our immunofluorescence analysis highlighted a notable presence of CD68-positive macrophages within the neointimal hyperplasia. This indicates an ongoing inflammatory response, which can contribute to adverse tissue remodeling and exacerbate the development of neointimal hyperplasia over time.

In contrast to ePTFE graft, the DAM grafts showed more favorable outcomes in terms of tissue integration and inflammatory response. Immunofluorescence staining revealed organized smooth muscle cell alignment in the middle layer and limited presence of CD68-positive macrophages within DAM grafts, indicating a stable and biocompatible integration into the vascular environment. This superior cellular response suggests that DAM grafts may facilitate a more natural healing process compared to synthetic counterparts, potentially reducing the risk of neointimal hyperplasia and thrombotic events over time. These findings underscore the potential clinical advantages of DAM grafts in enhancing vascular healing and maintaining long-term patency, positioning them as promising alternatives for vascular repair and reconstruction.

The findings from this study highlight the pressing need for advancements in graft materials that can effectively support endothelialization and mitigate inflammation. Traditional synthetic grafts, exemplified by ePTFE, often face challenges such as neointimal hyperplasia and impaired long-term patency due to inadequate reendothelialization and persistent inflammatory responses. The ability of DAM grafts to promote improved cellular organization and mitigate chronic inflammation may contribute to enhanced graft longevity. By supporting endothelial cell attachment and growth, DAM grafts offer a pathway to improve vascular healing processes, which are critical for long-term patency.

## 5. Conclusion

This study demonstrated the successful transplantation of DAM grafts in a pig for carotid artery replacement, achieving comparable outcomes and potential advantages to the conventional ePTFE grafts in terms of surgical feasibility, patency, and post-operative recovery. The DAM scaffold presents a versatile and adaptable solution for vascular grafting. Its preservation of ECM structure and mechanical properties following decellularization, along with the ability to customize mechanical stiffness by varying the layers, positions it as a highly promising candidate for clinical applications. Furthermore, the ePTFE grafts revealed significant proliferation in the neointimal layer, a typical response leading to luminal narrowing and graft failure. Notably, there was an absence of endothelial cells, crucial for preventing thrombosis and maintaining vascular function, and a notable presence of macrophages, indicating an ongoing inflammatory response. In contrast, DAM grafts showed organized SMC alignment, limited macrophage presence, and better tissue integration, indicating a natural healing process and reduced risks of neointimal hyperplasia and thrombotic events. These findings highlight the potential clinical advantages of DAM grafts for vascular repair and long-term patency. Future research should focus on strategies to further enhance endothelialization of DAM grafts and investigate the long-term outcomes in various vascular environments. Additionally, exploring the molecular mechanisms underlying the favorable response to DAM grafts could provide valuable insights for the development of next-generation vascular grafts. These studies will be crucial in establishing the clinical efficacy of DAM grafts and their potential to improve patient tailored outcomes in vascular surgery.

## Acknowledgements

Research reported in this publication was supported by the Advancing a Healthier Wisconsin (AHW) Endowment.

## Author contributions

Conceptualization: BW

Methodology: OA, PR, MD, AS, YZ, AK, XW, YC, BT, RW, LG, BW

Investigation: OA, PR, MD, AS, YZ, AK, XW, YC, BT, RW, LG, BW

Visualization: OA, BW

Supervision: BW

Writing: OA, BW

## References

1. Zaragoza, C., et al., Animal models of cardiovascular diseases. J Biomed Biotechnol, 2011. 2011: p. 497841.

2. Giannoukas, A.D., et al., Pre-bypass quality assessment of the long saphenous vein wall with ultrasound and histology. Eur J Vasc Endovasc Surg, 1997. 14(1): p. 37–40.

3. Leopold, J.A., et al., Diagnosis and Treatment of Right Heart Failure in Pulmonary Vascular Diseases: A National Heart, Lung, and Blood Institute Workshop. Circ Heart Fail, 2021. 14(6).

4. Shi, R. and S. Babu, Modern approaches and innovations in the diagnosis and treatment of peripheral vascular diseases. Front Biosci (Schol Ed), 2021. 13(2): p. 173–180.

5. Wu, Y.G., et al., Application of the Willis Covered Stent in the Treatment of Complex Vascular Diseases of the Internal Carotid Artery and Vertebral Artery: A Retrospective Single-Center Experience. Ther Clin Risk Manag, 2023. 19: p. 773–782.

6. Kim, E.S.H. and J.A. Beckman, Introduction to the Vascular Medicine Issue of Progress in Cardiovascular Diseases. Prog Cardiovasc Dis, 2018. 60(6): p. 565–566.

7. Kodama, T., [System biological medicine and treatment of vascular diseases]. Jpn J Antibiot, 2004. 57 Suppl A: p. 109–13.

8. Matsumoto, H., et al., Clinical results of 30 consecutive patients of carotid artery stenosis treated with CASPER stent placement: 1-year follow-up and in-stent findings on intravascular ultrasound examination immediately and 6 months after treatment. J Neurointerv Surg, 2023.

9. Megaly, M., et al., Outcomes of intravascular brachytherapy for recurrent drug-eluting in-stent restenosis. Catheter Cardiovasc Interv, 2021. 97(1): p. 32–38.

10. Kim, D.W., Breaking From the Past: Intravascular Stent Therapy in Pediatrics. JACC Cardiovasc Interv, 2016. 9(11): p. 1150–1.

11. Tara, S., et al., Vessel bioengineering. Circ J, 2014. 78(1): p. 12–9.

12. Beygui, F., et al., Long-term outcome after bare-metal or drug-eluting stenting for allograft coronary artery disease. J Heart Lung Transplant, 2010. 29(3): p. 316–22.

13. Elbaz, M., et al., Does stent design affect the long-term outcome after coronary stenting? Catheter Cardiovasc Interv, 2002. 56(3): p. 305–11.

14. Kastrati, A., D. Hall, and A. Schomig, Long-term outcome after coronary stenting. Curr Control Trials Cardiovasc Med, 2000. 1(1): p. 48–54.

15. Migliorini, A., et al., High residual platelet reactivity after clopidogrel loading and long-term clinical outcome after drug-eluting stenting for unprotected left main coronary disease. Circulation, 2009. 120(22): p. 2214–21.

16. Tiroch, K., et al., Impact of coronary anatomy and stenting technique on long-term outcome after drug-eluting stent implantation for unprotected left main coronary artery disease. JACC Cardiovasc Interv, 2014. 7(1): p. 29–36.

17. Aper, T., et al., Autologous blood vessels engineered from peripheral blood sample. Eur J Vasc Endovasc Surg, 2007. 33(1): p. 33–9.

18. Galego, S.J., et al., Blood flow study of arteriovenous grafts with homologous and autologous veins in canine femoral vessels. J Vasc Access, 2006. 7(1): p. 15–23.

19. Gavrilenko, A.V., et al., [Autologous vein bypass surgery in situ for reconstructive surgery of blood vessels of the leg]. Vestn Ross Akad Med Nauk, 1997(11): p. 39–42.

20. Hoenig, M.R., et al., Tissue-engineered blood vessels: alternative to autologous grafts? Arterioscler Thromb Vasc Biol, 2005. 25(6): p. 1128–34.

21. Desai, N.D., et al., A randomized comparison of radial-artery and saphenous-vein coronary bypass grafts. N Engl J Med, 2004. 351(22): p. 2302–9.

22. Klinkert, P., et al., Vein versus polytetrafluoroethylene in above-knee femoropopliteal bypass grafting: five-year results of a randomized controlled trial. J Vasc Surg, 2003. 37(1): p. 149–55.

23. Zhuang, Y., et al., Challenges and strategies for in situ endothelialization and long-term lumen patency of vascular grafts. Bioact Mater, 2021. 6(6): p. 1791–1809.

24. Tatterton, M., et al., The use of antithrombotic therapies in reducing synthetic small-diameter vascular graft thrombosis. Vasc Endovascular Surg, 2012. 46(3): p. 212–22.

25. Fellows, E., Vascular graft research: endothelial cell seeding of synthetic bypass grafts. J Vasc Nurs, 1991. 9(2): p. 12–4.

26. Zylke, J.W., Synthetic vascular graft trials start; endothelialization seen as possible. JAMA, 1990. 264(20): p. 2607–8.

27. Abdallah, O.I., D. Clark, and M.A. Essibayi, Surgical Reconstruction of a Traumatic Superior Sagittal Sinus Injury Using Synthetic Vascular Graft in a Resource-Limited Civilian Field Hospital During the Syrian Civil War. World Neurosurg, 2022. 159: p. 126–129.

28. Michael, P.L., et al., Synthetic vascular graft with spatially distinct architecture for rapid biomimetic cell organisation in a perfusion bioreactor. Biomed Mater, 2022. 17(4).

29. Campbell, G.R. and J.H. Campbell, Development of tissue engineered vascular grafts. Curr Pharm Biotechnol, 2007. 8(1): p. 43–50.

30. Kuang, H., et al., Construction and performance evaluation of Hep/silk-PLCL composite nanofiber small-caliber artificial blood vessel graft. Biomaterials, 2020. 259: p. 120288.

31. Wang, B., et al., Developing small-diameter vascular grafts with human amniotic membrane: long-term evaluation of transplantation outcomes in a small animal model. Biofabrication, 2023. 15(2).

32. Dhaliwal, G. and D. Mukherjee, Peripheral arterial disease: Epidemiology, natural history, diagnosis and treatment. Int J Angiol, 2007. 16(2): p. 36–44.

33. Virani, S.S., et al., Heart Disease and Stroke Statistics-2021 Update: A Report From the American Heart Association. Circulation, 2021. 143(8): p. e254–e743.

34. Brahmbhatt, A. and S. Misra, The Biology of Hemodialysis Vascular Access Failure. Semin Intervent Radiol, 2016. 33(1): p. 15–20.

35. Li, W., et al., Polymer-integrated amnion scaffold significantly improves cleft palate repair. Acta Biomater, 2019.

36. Wang, B., et al., Developing small-diameter vascular grafts with human amniotic membrane: long-term evaluation of transplantation outcomes in a small animal model. Biofabrication, 2023.

37. Li, W., et al., Investigating the Potential of Amnion-Based Scaffolds as a Barrier Membrane for Guided Bone Regeneration. Langmuir, 2015. 31(31): p. 8642–53.

38. Wang, B., W. Li, and J. Harrison, An Evaluation of Wound Healing Efficacy of a Film Dressing Made from Polymer-integrated Amnion Membrane. Organogenesis, 2020. 16(4): p. 126–136.

